# Single-cell transcriptomics of *Encephalitozoon intestinalis* infection in Caco-2 and Vero cells

**DOI:** 10.64898/2026.09.18.752679

**Authors:** Noelle V. Antao, Gira Bhabha, Damian C. Ekiert

## Abstract

Microsporidia are single-celled, obligate intracellular parasites closely related to fungi. One of the most common human-infecting microsporidian species, *Encephalitozoon intestinalis*, is an enteric parasite that invades and replicates within the epithelial lining of the small intestine. Two epithelial cell lines, Caco-2 and Vero cells, are routinely used as *in vitro* models of *E. intestinalis* infection. Here, we use scRNA-seq to investigate the host response to *E. intestinalis* infection and the transcriptional dynamics of parasite development in both Caco-2 and Vero cell lines. We observe that most host cells have no direct transcriptional response to parasite invasion or intracellular replication, suggesting that cells fail to detect and/or fail to respond to direct *E. intestinalis* infection, consistent with previous single-cell transcriptomics of *E. intestinalis*-infected macrophages. Our data suggest that a small subset of host cells detect intracellular parasites late in the life cycle, and signal to neighboring cells, leading to broad activation of a TNF-α response. Analysis of parasite transcripts reveals a developmental program that correlates with the morphological stages of the parasite life cycle. Together our data provide insights into a shared host transcriptional response to *E. intestinalis* infection and the datasets presented are a valuable resource for future studies.

## Introduction

Microsporidia are an early-diverging group of fungal parasites that infect most animals including humans. ~17 microsporidian species are known to infect humans [1] resulting in human microsporidian infections (microsporidiosis) that have diverse clinical manifestations. *Encephalitozoon intestinalis* is one of the most common human-infecting species [2], and typically manifests as an enteric infection in which the parasites invade and replicate within the mucosal epithelial lining of the small intestine, and may then be disseminated to other organs. Such disseminated infections can be severe and even fatal in immunocompromised individuals, such as AIDS patients, organ transplant recipients, and those with autoimmune disorders [3–5]. Like all microsporidia, *E. intestinalis* is an obligate intracellular parasite, and the only life cycle stage that can survive outside the host is the dormant spore stage. Infection begins when a spore germinates, releasing a long invasion organelle called the polar tube, which is thought to facilitate parasite entry, either by direct injection into the cytoplasm or through a phagocytosis-like mechanism [6]. Once inside the host, *E. intestinalis* develops within a vacuole called the parasitophorous vacuole, and transitions through a series of four stages: proliferative cells undergoing binary fission; sporonts, a stage in which the spore coat begins to develop; sporoblasts, a post-replicative stage during which the polar tube develops; and spores, the transmissible form of the parasite [7–9]. Mature spores are released from the host into the environment, where they remain dormant until infection of a new host.

Previous bulk RNA sequencing and single-cell RNA sequencing (scRNA-seq) studies have provided insights into the transcriptional dynamics of *E. intestinalis* development and the host response to infection [10–14]. Bulk RNA sequencing analysis of the host response to *E. intestinalis* infection in Caco-2 cells, a human colon carcinoma cell line, showed impacts on host mitochondrial function, cellular metabolism and membrane trafficking [13]. scRNA-seq analysis of human macrophages infected with *E. intestinalis* revealed that most cells fail to detect or respond transcriptionally to intracellular parasites [15]. However, a small population does respond, and may signal to neighboring cells, resulting in a transcriptional response in nearby uninfected and infected cells. Analysis of parasite transcripts from the same datasets established the overall transcriptional landscape for *E. intestinalis* across its lifecycle in macrophages. Early developmental stages express housekeeping genes, intermediate developmental stages express genes involved in protein synthesis and long chain fatty acid synthesis, and late developmental stages express genes necessary for spore coat synthesis and polar tube development.

To better understand the similarities and differences in the host response to *E. intestinalis* infection in different cell types, and how the transcriptional dynamics of parasite development may enable adaptation to distinct cell types, we performed scRNA-seq in Caco-2 cells and Vero cells infected with *E. intestinalis*, and analyzed both host and parasite transcriptomes. Caco-2 cells are a colon carcinoma cell line that is used as a model of the intestinal epithelial barrier [16], and Vero cells are African green monkey kidney epithelial cells, and are characterized by a deletion in type I interferon genes. These cell lines are commonly used to study *E. intestinalis in vitro*. Therefore, obtaining information on the transcriptional landscape of both the host and parasite in these cell lines would be a valuable resource to aid the interpretation of future *E. intestinalis* host-pathogen studies.

## Results

### scRNA-seq of Caco-2 and Vero cells infected with E. intestinalis

To identify which infection time points to select for scRNA-seq analysis, we infected Caco-2 and Vero cells with *E. intestinalis* and monitored infection by light microscopy, from 12 to 42 hours post infection (hpi). Cells were stained with a fluorescent DNA dye (DRAQ5), which stains all stages of parasite development, as well as the host nucleus; and with a fluorescent chitin dye (calcofluor white), which stains the spore coat, allowing us to monitor the development of spores. To identify early infection events, we used an RNA fluorescence in situ hybridization (FISH) probe against *E. intestinalis* 16S rRNA [9]. At 12 hpi, ~40% of host cells were infected for both the Caco-2 and Vero cell lines (Figure 1A-C). At this time point, either a single FISH punctum or a chain of two FISH puncta were observed in infected cells, representing early infection stages (Figure 1A). From 12 hours to 24 hours, infection rates increased, with ~54% of Caco-2 cells and 60% of Vero cells being infected (Figure 1A-C). By 18 and 24 hpi, longer chains of FISH puncta were observed (Figure 1A). After peaking at 24 hpi, infection rates drop slightly at later time points, to 40% in Caco-2 cells and 54% in Vero cells by 42 hpi. Intracellular spores were first detected at 24 hpi in both Caco-2 and Vero cells, with ~5% of infected cells in both cell lines containing chitin positive spores (Figure 1B-C). In both cell lines, from 24 hours to 42 hpi, the number of infected cells with chitin positive spores increased (Figure 1B-C). To investigate 1) the host transcriptional response to *E. intestinalis* infection and 2) the dynamics of *E. intestinalis* transcription during parasite development, and how it may differ between cell types, we proceeded with scRNA-seq of *E. intestinalis*-infected Caco-2 and Vero cells at 12 hpi, 18 hpi, 24 hpi, 36 hpi and 42 hpi, along with uninfected control cells (Figure 1D).

**Figure 1.**
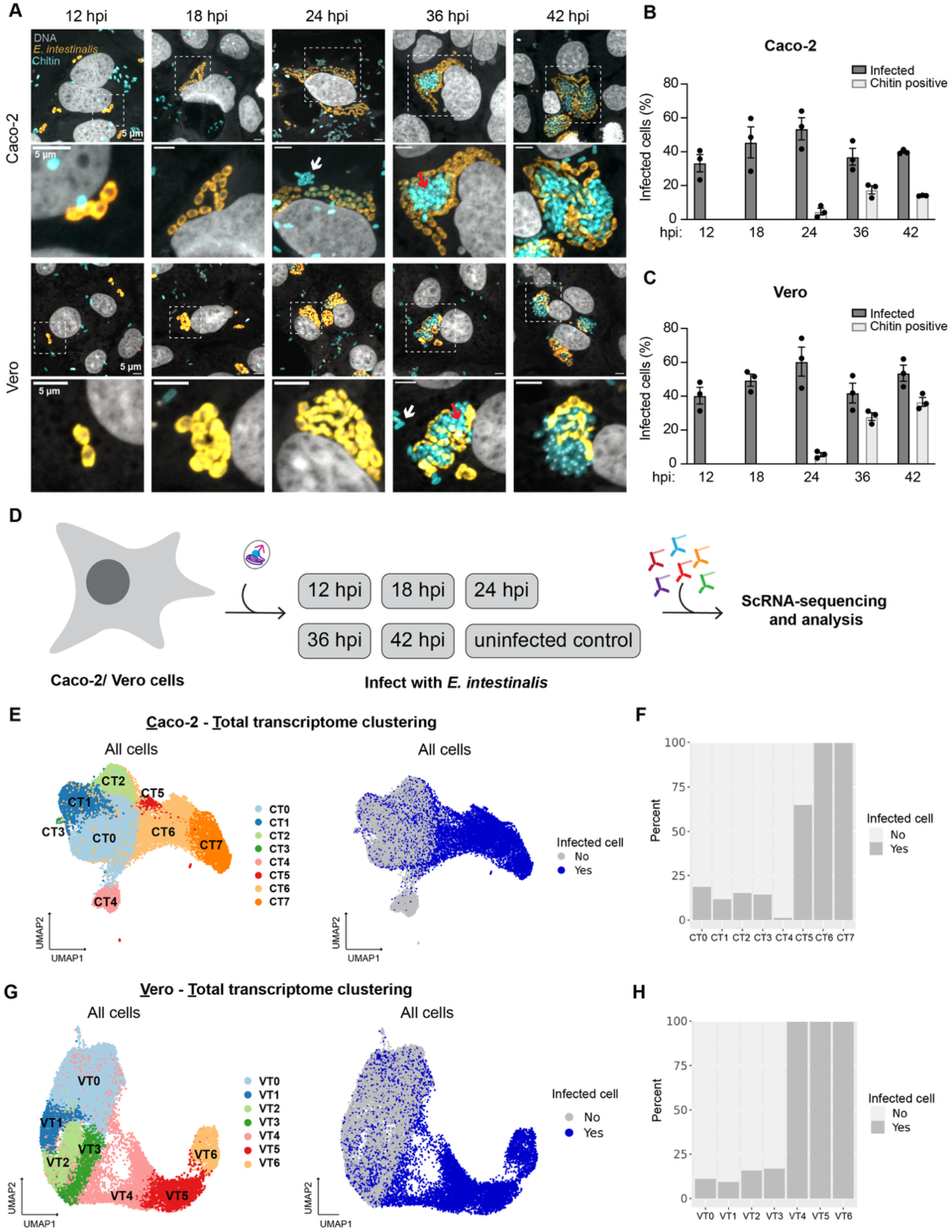
Infection kinetics of *E. intestinalis* in Caco-2 and Vero cells. **A.** Representative micrographs of Caco-2 cells and Vero cells infected with *E. intestinalis* for 12, 18, 24, 36 and 42 hpi. Cells are stained for DNA (DRAQ5, gray), chitin (calcofluor white, cyan) and *E. intestinalis* (RNA FISH probe, orange). White arrows indicate extracellular spores and red arrows indicate intracellular spores. **B**. Quantification of infected Caco-2 cells at 12, 18, 24, 36 and 42 hpi. Mean ± SEM are from three biological replicates, n= 1,917 cells. **C**. Quantification of infected Vero cells at 12, 18, 24, 36 and 42 hpi. Mean ± SEM are from three biological replicates, n= 1,889 cells. **D**. Schematic of the experimental workflow for scRNA-seq experiments. Caco-2 or Vero cells were infected with *E. intestinalis* for 12, 18, 24, 36 or 42 hours alongside uninfected control cells. Cells were then hashed and subjected to the scRNA-seq workflow. **E**. UMAP plot of host and parasite transcripts (left) and colored by uninfected cells and infected cells (right) across all time points for the Caco-2 infected cell dataset. **F**. Quantification of the percentage of infected cells in each cluster of the Caco-2 infected cell dataset. **G**. UMAP plot of host and parasite transcripts (left) and colored by uninfected cells and infected cells (right) across all time points for the Vero infected cell dataset. **H**. Quantification of the percentage of infected cells in each cluster of the Vero infected cell dataset.

After quality control and filtering of cells (see Methods), we analyzed 22,748 Caco-2 cells and 24,543 Vero cells across all infection timepoints and uninfected controls (Supplementary Figure 1A and B). To determine which host cells were infected, we calculated the percentage of *E. intestinalis* transcripts detected in each cell and assessed thresholds for scoring a cell as “infected” between 0.2% and 80% microsporidia transcripts (Supplementary Figure 1C-F). A 0.2% cut-off for microsporidia transcripts maximizes the recovery of cells originating from experimentally infected samples that are scored as infected, while minimizing the number of false positives (cells originating from the uninfected control samples that are scored as infected; Supplementary Figure 1D&F; see Methods). Thus, we proceeded with a threshold of 0.2% *E. intestinalis* transcripts to define a cell as infected. Following data normalization and unsupervised cell clustering, Caco-2 cells were partitioned into 8 clusters (CT0-CT7, where “CT” denotes “Caco-2 - Total transcriptome”) and Vero cells were partitioned into 7 clusters (VT0-VT6, where “VT” denotes “<u>V</u>ero-<u>T</u>otal transcriptome”), and visualized using a Uniform Manifold Approximation and Projection (UMAP) (Figure 1E&G). However, differential gene expression (DGE) analysis of the combined host and parasite transcripts revealed that cells were being clustered primarily based on the presence or absence of *E. intestinalis* transcripts, and not on differential transcription in host cells (Figure 1F&H and Supplementary Figure 1G-H). Therefore, to focus on changes to the host cell transcriptome in response to infection, while also gaining insights into the parasite developmental program in each cell line, we analyzed the host and parasite transcripts separately, as previously described in scRNA-seq analysis of *E. intestinalis*-infected macrophages [15].

### Transcriptional dynamics of E. intestinalis in Caco-2 and Vero cells

To examine the dynamics of *E. intestinalis* transcription during parasite development, we analyzed the parasite transcripts only from both the Caco-2 and Vero infected cell datasets. As many *E. intestinalis* parasites develop within a single host cell, and it is the host cell that is being partitioned in the scRNA-seq experiment, this analysis represents the transcriptome of multiple developing parasites present in each infected host cell, which may be developing somewhat asynchronously. After quality control and filtering, 10,915 infected Caco-2 cells and 11,192 infected Vero cells were analyzed across all timepoints, with each infected cell containing at least 0.2% microsporidian transcripts (Supplementary Figure 2C).

Following data normalization and unsupervised clustering, infected cells were partitioned into 6 distinct clusters in the Caco-2 infected cell dataset (P0-P5, where P denotes Parasite) (Figure 2A) and similarly, 6 distinct clusters in the Vero infected cell dataset (P0-P5) (Figure 3A). To investigate the trajectory of parasite development through these clusters, we set out to identify which clusters may correspond to earlier stages of infection and which may correspond to later stages of infection. To this end, we used the percentage of *E. intestinalis* transcripts as a proxy for parasite burden in each cell, assuming that parasite burden roughly correlates with infection stage. We also assessed the distribution of infected cells across clusters as a function of infection time point. We observed that the proportion of parasite transcripts varied across clusters, with the lowest proportion in cluster P0, and increasing in abundance in clusters P1, P2, P3, P4 and P5 in both Caco-2 and Vero cells (Figure 2B and 3B). This trend indicates that parasite burden is lowest in P0 and highest in P5, suggesting that P0 represents the earliest stage of infection and that P5 represents the latest stage of infection. In line with this interpretation, nearly all infected cells from the 12 hpi samples are found in clusters P0, P1, and P2 (Supplementary Figure 2A-B and 2D-E), while clusters P4 and P5 are nearly absent. Some cells are observed in cluster P3 at 12 hpi in both cell lines, and P3 expands further by 18 hpi, along with the appearance of cells in P4. By 24 hpi, all 6 parasite clusters are well populated, including the final cluster, P5 (Supplementary Figure 2A-B and 2D-E). Based on the above, we propose that parasite development proceeds in numerical order from P0 (immediately after entry) to P5 (the latest stages of infection) and that the overall trajectory of parasite development is very similar in both cell lines (Supplementary Figure 2A and 2D). We tentatively assign these transcriptional states as follows, and will discuss the rationale behind these assignments in the next section: post-invasion/pre-proliferative (P0/P1/P2); proliferative stage (P3); sporont stage (P4); and sporoblast (P5).

**Figure 2.**
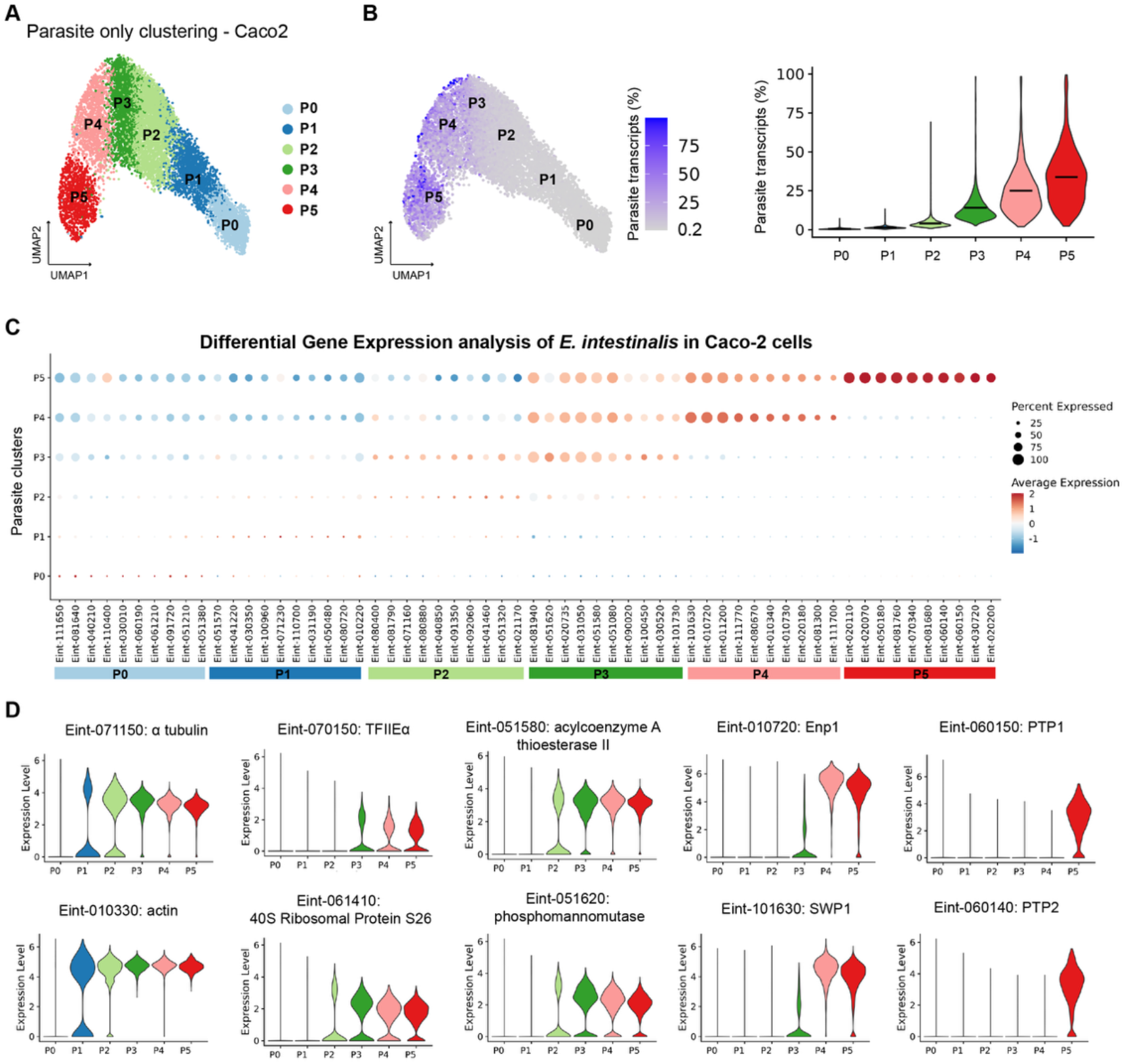
Gene expression dynamics of *E. intestinalis* during parasite development in Caco-2 cells. A. UMAP plot of parasite-only transcripts where each cluster represents a parasite developmental stage (P0-P5). **B.** UMAP plot of parasite-only transcripts colored by the percentage of microsporidia transcripts (left) and corresponding quantification of the percentage of parasite transcripts per cluster (right). **C**. Dotplot of the top 10 differentially expressed genes in each parasite cluster. The X-axis shows the genes and the Y-axis shows the parasite clusters. The color represents the average gene expression level across the cells in each parasite cluster in which it is expressed and the size indicates the percentage of cells in each parasite cluster expressing that gene. **D**. Violin plots showing the expression levels for the top differentially expressed genes across parasite clusters.

### Differential gene expression analysis of E. intestinalis development in Caco-2 and Vero cells

Levels of *E. intestinalis* RNA are very low in P0 and P1, and increase modestly in P2 (Fig. 2B and 3B), which may represent initial transcription in the invading parasites, before the first cell division. By light microscopy, most parasite cells have yet to divide by 12 hpi, though some have begun the first division, and most will divide several times by 18 hpi. Clusters P0-P2 are well populated by 12 hpi, and cells are just beginning to enter P3, which will expand fully by 18 hpi. Thus, the initiation of cell division appears to coincide with the formation and expansion of cluster P3. DGE analysis of clusters P0-P3 revealed that the magnitude of transcriptional differences in these early parasite clusters is modest in Caco-2 and Vero cells, ranging from 1.5-fold to 4-fold, similar to previous analyses of *E. intestinalis* in macrophages [15]. In clusters P0-P2, we observe an upregulation in genes necessary for transcription and translation including transcription initiation factors, ribosomal proteins and cytoskeletal proteins like actin and tubulin (Figure 2C-D and 3C-D, Supplementary Table 1&2). In cluster P3 we observe an upregulation of genes involved in lipid metabolism and glycosylation.

**Figure 3.**
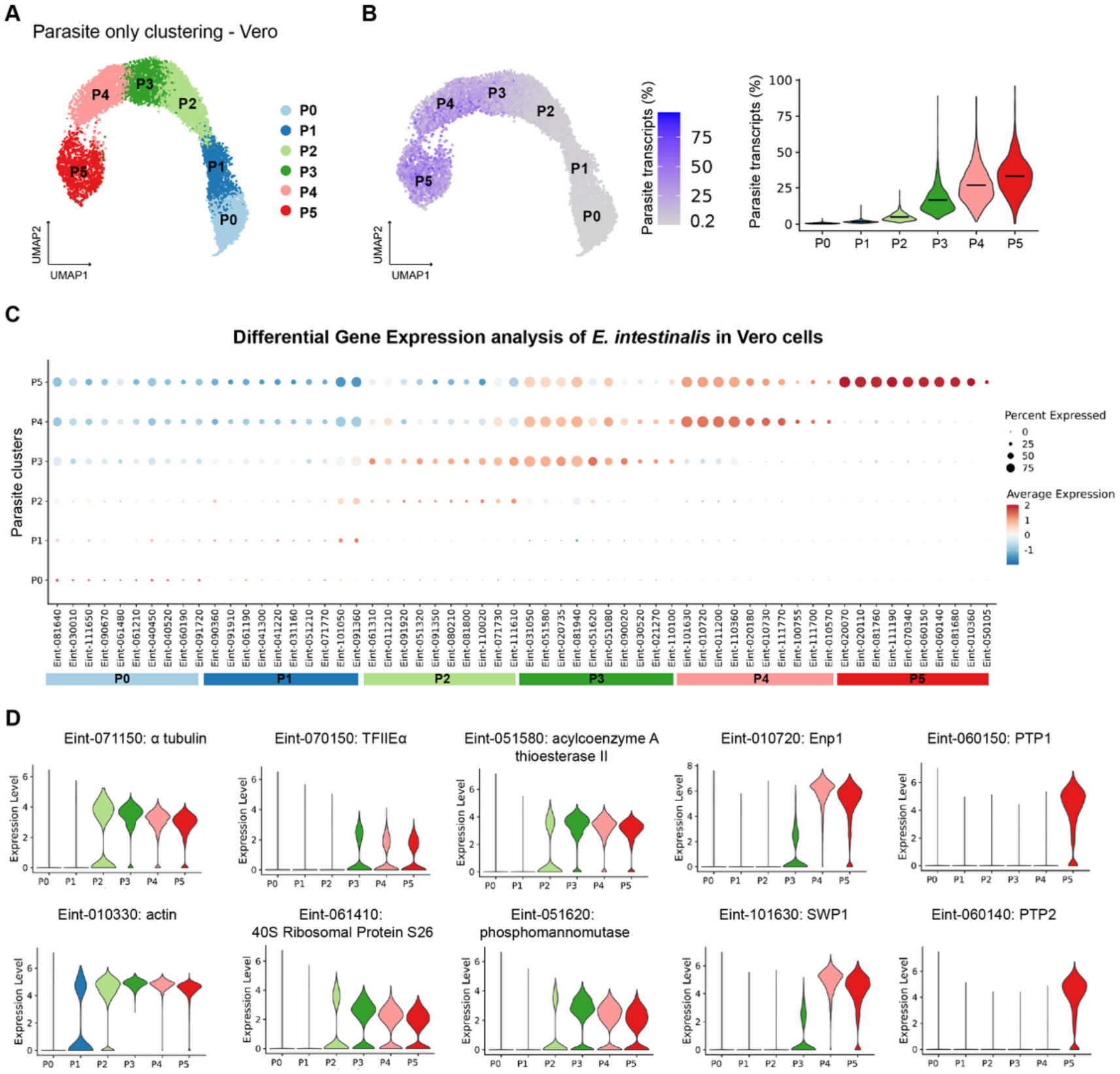
Gene expression dynamics of *E. intestinalis* during parasite development in Vero cells. **A.** UMAP plot of parasite-only transcripts where each cluster represents a parasite developmental stage (P0-P5). **B**. UMAP plot of parasite-only transcripts colored by the percentage of microsporidia transcripts (left) and quantification of the percentage of parasite transcripts per cluster (right). **C**. Dotplot of the top 10 differentially expressed genes in each parasite cluster. The X-axis shows the genes and the Y-axis shows the parasite clusters. The color represents the average gene expression level across the cells in each parasite cluster in which it is expressed and the size indicates the percentage of cells in each parasite cluster expressing that gene. **D**. Violin plots showing the expression levels for the top differentially expressed genes across parasite clusters.

Consequently, we propose that the P0, P1, and P2 states, which are transcriptionally similar to one another, likely represent the invading sporoplasm and unicellular states as parasite cells initiate transcription and begin to produce the basic building blocks of the growing cell. Following this growth initiation phase, P3 represents the major proliferative state of *E. intestinalis*, wherein the parasites undergo multiple rounds of genome replication and binary fission. Levels of parasite RNA increase substantially in P3 and the width of the distribution also increases relative to earlier stages (Fig. 2B and 3B), consistent with an overall increase in parasite burden and infected host cells with variable numbers of intracellular parasites.

Clusters P4 and P5 mark a shift away from general parasite proliferation, towards differentiation into transmissible spores. The magnitude of the transcriptional changes in clusters P4 and P5 is also much larger, more than 10-fold for many genes. For cluster P4, the most differentially expressed genes include genes necessary for the synthesis of the two layers of the spore coat: endospore protein 1 (EnP1), chitin synthase, and spore wall protein 1 (SWP1) (Figure 2C-D and 3C-D). The hallmark of the transition from the proliferative stages to the sporont stage is the appearance of a thin layer of spore coat around the parasites. Thus, we assign the P4 state as corresponding to the sporont stage of parasite development, as the transcriptional profile of P4 is in good agreement with morphological observations, and a similar transcriptional state was previously proposed to represent sporonts in *E. intestinalis*-infected macrophages [15]. Levels of parasite RNA continue to increase in P4 relative to earlier stages (Fig. 2B and 3B), likely reflecting a combination of the asynchronous parasite stages within each host cell and some continued division of proliferative forms in the same cell. Finally, in cluster P5 we observe the largest transcriptional changes, with up to a 64-fold upregulation in the expression of several specialized genes, including polar tube protein 1 (PTP1) and polar tube protein 2 (PTP2), core components of the microsporidian polar tube (Figure 2C-D and 3C-D). The initiation of polar tube assembly marks the transition from the sporont to the sporoblast stage, and thus, the upregulation of polar tube protein expression in this cluster supports the assignment of P5 as the sporoblast stage. In addition to polar tube development, the sporoblast stage is also marked by continued development and thickening of the spore coat, ultimately resulting in mature spores. In our datasets, we do not expect to capture transcripts from fully mature *E. intestinalis* spores, as the thick spore coat precludes efficient RNA release by the detergent-based lysis methods used in scRNA-seq library construction.

The overall *E. intestinalis* transcriptional program appears to be globally similar between the host cell types investigated thus far by scRNA-seq, including Caco-2, Vero, and monocyte-derived human macrophages [15]. To assess whether there are noteworthy host cell-specific differences in parasite gene expression, we compared the top 50 differentially expressed genes in comparable clusters across all three cell types (P0, P4 and P5) (Supplementary Figure 3A-C). Due to differences in clustering between the Caco-2/Vero and macrophage [15] analyses, comparison of clusters P1-P3 was only performed for the Caco-2 and Vero infected cell datasets (Supplementary Figure 3D and see Methods). The data were generally well correlated across the three datasets. However, we observed some differences, with 12 genes being differentially expressed across the different host cell lines. For the earliest parasite developmental stage (P0), we found Eint_060230 (DNA directed RNA polymerase) has ~8 fold higher expression in Caco-2 cells versus macrophages and ~1.8 fold higher expression in Caco-2 compared to Vero cells. Another gene, Eint_051080 (hypothetical protein), has ~6 fold higher expression in macrophages compared to Vero cells and 1.4 fold higher expression in Caco-2 cells compared to Vero cells (Supplementary Figure 3A). For the later developmental stages, differentially expressed genes in cluster P4 were tightly correlated between datasets, with no significant difference in gene expression (Supplementary Figure 3B). However, in cluster P5, we observe two genes that are differentially expressed in the different host cells. Eint_050105 (hypothetical protein) is consistently expressed higher in Vero cells when compared to Caco-2 cells (1.8 fold) and macrophages (14 fold), and Eint_010620 (Leptin receptor) has 3-fold higher expression in Caco-2 cells when compared to Vero cells but is similarly expressed in Macrophages (Supplementary Figure 3C). Taken together, these data suggest that, overall, parasite developmental trajectory is well correlated across the different cell lines, but some host-specific differences may fine-tune parasite development in different hosts.

### Transcriptional response to E. intestinalis infection in Caco-2 host cells

We next focused on the host transcriptional response to *E. intestinalis* infection in Caco-2 cells. Following data normalization and unsupervised cell clustering, cells were partitioned into 8 clusters in the Caco-2 infected cell dataset (C0-C7, where “C” denotes “Caco-2”) (Figure 4A). Analysis of clusters in Caco-2 cells at different infection time points revealed that the distribution of cells across the clusters changes substantially over the course of infection (Figure 4C-D). Relative to the uninfected control, the most populated clusters are depleted of cells over the course of infection, with the populations of clusters C0, C1, and C3 decreasing 4- to 9-fold by 42 hpi. In contrast, clusters C2 and C5 have the opposite trend, expanding more than 10-fold and together accounting for ~81% of cells by 42 hpi (Figure 4C-D). We observed that most clusters consist of a mixture of ~40-75% infected cells that are interspersed with uninfected cells (Figure 4A-B), indicating that the transcriptomes of infected host cells are generally indistinguishable from uninfected cells. In total, the majority of infected cells (~97%) fall into this category, suggesting that most host cells do not appreciably alter transcription in direct response to infection. This could be due to a failure of the host cell to detect *E. intestinalis* and mount a response, or due to parasite inhibition of anti-microsporidian host responses. To understand what differentiates the 8 host cell clusters from one another, we performed DGE analysis and identified the top marker genes that are unique to each of the cell clusters (Figure 4E). Three clusters are of particular interest, and are discussed in more detail below: 1) C5, which is induced by infection and has a very high infection rate, 2) C2, which is induced by infection and accounts for the vast majority of cells analyzed at late infection time points, and 3) C3, which has a low infection rate.

**Figure 4.**
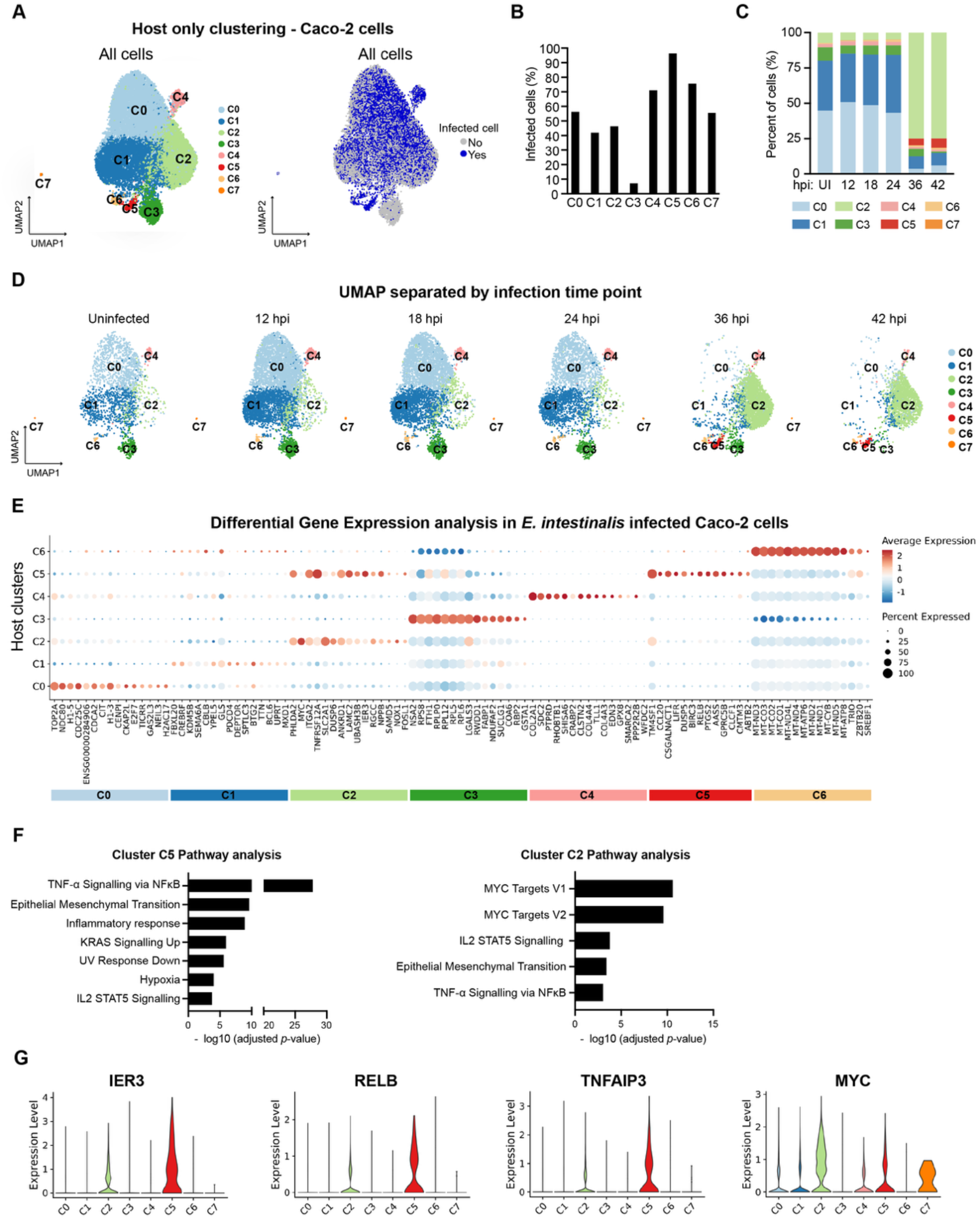
Analysis of host transcriptional response to *E. intestinalis* infection in Caco-2 cells. **A.** UMAP plot of host-only transcripts for all cells (left) and colored by uninfected cells and infected cells (right) across all time points. **B**. Quantification of the percentage of infected cells in each host cluster (n=22,748 cells). **C**. Quantification of the proportion of cells per cluster at 12, 18, 24, 36 and 42 hpi along with uninfected (UI) control cells. **D**. UMAP plot of host-only transcripts separated by infection time point. **E**. Dotplot of the top 15 differentially expressed genes in each host cluster. The X-axis shows the genes and the Y-axis shows the host clusters. Cluster C7 was excluded from DGE analysis as it only contained 27 cells. The red-blue color range represents the average gene expression level across the cells in each host cluster in which it is expressed and the size indicates the percentage of cells in each host cluster expressing that gene. **F**. Bar plots showing −log_10_ adjusted *p*-value from gene set enrichment analysis of biological pathways on host clusters C5 and C2 using the hallmark gene set in the Human Molecular Signatures Database (MSigDB) [31,32]. **G**. Violin plots showing the expression levels for the top differentially expressed genes in host clusters C5 and C2.

Cluster C5 is unique in that it consists almost entirely of infected cells (96% infected, Figure 4B) and is induced in the later stages of infection (only cells from the 36 hpi and 42 hpi samples contribute to cluster C5; Supplementary Figure 1A). One possibility is that cells in this cluster have altered host transcription in direct response to infection. Notably, cluster C5 represents just 2.7% of all infected cells, which may indicate that only a very small subset of infected cells activate this response, or that the response may be very short-lived. Relative to other host cell clusters, C5 cells have higher levels of *E. intestinalis* RNA (Supplementary Figure 4A), indicating that C5 cells have higher parasite burden, and could represent later stages of the infection cycle. To investigate this further, we assessed the distribution of parasite transcriptional stages in each host cluster. For cluster C5, ~87% of parasites were in the latest stages (parasite developmental stages P4 and P5), while late stage parasites only accounted for ~11-40% of cells in other clusters (Supplementary Figure 4D). This suggests that a transcriptional response to intracellular parasites, represented by the cells in cluster C5, is usually triggered very late in infection. To better understand the transcriptional changes occurring in cluster C5, we performed gene set enrichment analysis (GSEA) on the differentially expressed genes identified in this cluster (Supplementary Table 3). This analysis revealed a robust induction of genes in pathways including TNF-α signaling via NF-κB, inflammatory response and IL2-STAT5 signaling (Figure 4E-G). This included genes like *RELB* and *TNFAIP3*, that regulate the host immune response through the NF-κB signalling pathway and *IER3*, an immediate-early response gene induced in response to cellular stress, including viral infection [17–19]. *RELB* is a component of the non-canonical NF-κB pathway, which is the target of some pathogens including RNA viruses and bacteria [20–22] and has a role in immune regulation and inflammatory disease. Conversely, *TNFAIP3* is a negative regulator of NF-κB signaling that prevents excessive inflammation and cellular damage in response to microbial products and viral infection [23,24]. Together these data suggest that the cells in cluster C5 usually contain parasites in the late stages of the lifecycle, have sensed infection, and triggered an inflammatory response in order to limit infection.

A second cluster, C2, was also induced by infection, and accounts for most of the cells at later infection time points in Caco-2 cells. While this cluster accounts for just ~5-7% of cells from the 12 hpi, 18 hpi, 24 hpi, and uninfected control samples; by 36 and 42 hpi, cluster C2 has expanded to encompass ~75% of all cells (Figure 4C-D). In parallel, the other major clusters shrink substantially, suggesting that infection induces cells from other clusters to alter transcription to the C2 state. However, in contrast to the high infection rate of cluster C5, only ~46% of cells in C2 are infected (Figure 4A-B), suggesting that something other than direct infection is causing both infected and uninfected cells to adopt the C2 transcriptional state. This shift to the C2 state at later infection time points may reflect cell-cell signaling downstream of infection that is acting on most cells in the culture. GSEA of differentially expressed genes in this cluster reveals a signature related to that of cluster C5, with induction of genes in the signaling pathways such as TNF-α signaling via NF-κB, IL2-STAT5 signaling, and Epithelial-Mesenchymal Transition (Figure 4E-F). As in cluster C5, genes such as *IER3, RELB* and *TNFAIP3* were also upregulated in cluster C2 (Figure 4G). Cluster C2 also exhibited robust induction of MYC targets (Figure 4E-F). But in contrast to C5, which represents cells in late stages of infection, the infected cells in C2 contain lower levels of *E. intestinalis* mRNA and the distribution of parasite transcriptional states skews towards earlier, proliferative stages (only 44% late stages; Supplementary Figure 4D). One possibility is that cells in C2 could be responding to a signal generated between 24 and 36 hpi that acts on the majority of the cells in the population, both infected and uninfected. A plausible source of this signal is the cells in cluster C5, which appear to have sensed intracellular parasites and may be secreting TNF-α and other factors in response. Alternatively, infected cells in cluster C2 may represent a transcriptional state just prior to C5, during which parasites are transitioning from a replicative stage to a spore development stage that is sensed by the host, triggering a stronger inflammatory response.

While at least 40% of cells in most clusters are infected with *E. intestinalis*, the infection rate of cluster C3 was significantly lower: only ~7% of cells were infected (Figure 4B). C3 is a smaller cluster, representing just ~6% of cells in the Caco-2 infected cell dataset and is present at all infection time points. One possible explanation for the low infection rate of cluster C3 is that cells in C3 are resistant to infection. In this case, we might expect that cells in C3 express factors that restrict parasite entry or replication. A second possibility is that only uninfected cells can access or maintain a C3-like transcriptional state. For example, since *E. intestinalis* infection has been proposed to lead to cell cycle arrest [25], we would expect cells actively proceeding through the cell cycle to be largely uninfected. We performed DGE analysis followed by GSEA on genes upregulated in cluster C3 relative to all other clusters (Figure 4E), to assess whether transcriptional changes may support or refute either possibility. We found upregulation of genes involved in oxidative phosphorylation, MYC targets, fatty acid metabolism, and DNA repair (Supplementary Figure 4C), but these signatures do not immediately explain why C3 has a low level of infection. If cells in C3 were truly resistant to infection, one might expect any infected cells to primarily contain early stage parasites, and that development to later stages would be blocked. We observe all parasite developmental stages in cluster C3, suggesting that this is not the case (Supplementary Figure 4D). Together, these data are consistent with cluster C3 representing a transcriptional state that only uninfected cells can access, but future experiments will be needed to resolve the basis for this transcriptional state

### Transcriptional response to E. intestinalis infection in Vero host cells

We next investigated the host response to infection in Vero cells. Cells were partitioned into 7 clusters (V0-V6, where “V” denotes “Vero”) following data normalization and unsupervised cell clustering (Figure 5A). UMAP visualization of cell clusters in Vero cells revealed that clusters V0-V4 and V6 are present in uninfected samples and their populations remain fairly stable across all infection timepoints (Figure 5C-D and Supplementary Figure 1B). However, cluster V5 is induced upon infection, with 99% of the cells in this cluster originating from infected samples (Figure 5C-D and Supplementary Figure 1B). Cluster V5 was enriched for infected cells, with 84% of cells in this cluster being infected, compared with ~40-60% infection rates in other clusters (Figure 5B and Supplementary Figure 4B). Thus, as in Caco-2 cells, and as previously observed in human macrophages [13,15], most infected cells are indistinguishable from neighboring uninfected cells based on the host transcriptome. Only V5 stands out from the other clusters and appears to be analogous to cluster C5 from Caco-2 cells, as it is induced by infection and the majority of cells from this cluster are infected. We therefore focused our analysis on cluster V5. As in Caco-2 cluster C5, the parasites in cluster V5 are transcriptionally characteristic of later stage infections, suggesting that infected cells usually adopt the V5 transcriptional state only late in infection, when parasite burden is likely quite high (Supplementary Figure 4E). DGE analysis and subsequent GSEA of upregulated genes in cluster V5 revealed enrichment of genes involved in TNF-α signalling via NF-κB and inflammatory response pathways (Figure 5E-F and Supplementary Table 4) including significant upregulation in genes like *IL1A, TNFAIP3* and *NFKBIA* (Figure 5G). While *IL1A* is a proinflammatory cytokine that initiates and amplifies inflammatory responses [26,27], *TNFAIP3* and *NFKBIA* serve as negative regulators of inflammation and limit NF-κB signalling. We also observed enrichment of genes involved in apoptosis signaling pathways (Figure 5F), which may reflect the release of late stage spores from the host by cell lysis. Together these responses suggest that similar to cluster C5 in Caco-2 cells, the Vero cells in cluster V5 represent late stages of infection, wherein the host cells have sensed infection and are mounting an inflammatory response. In contrast to the large redistribution of cells between Caco-2 clusters at the later infection time points (with most cells adopting a C2-like transcriptional state and a concomitant depletion of cells from other clusters), in Vero cells, the clusters are largely stable across the infection time course. This may be due to the inability of Vero cells to produce type I interferons, or some other signalling molecules, which induce the population-wide transcriptional changes observed in Caco-2 cells at later infection timepoints, as well as in *E. intestinalis*-infected macrophages from some donors [15].

**Figure 5.**
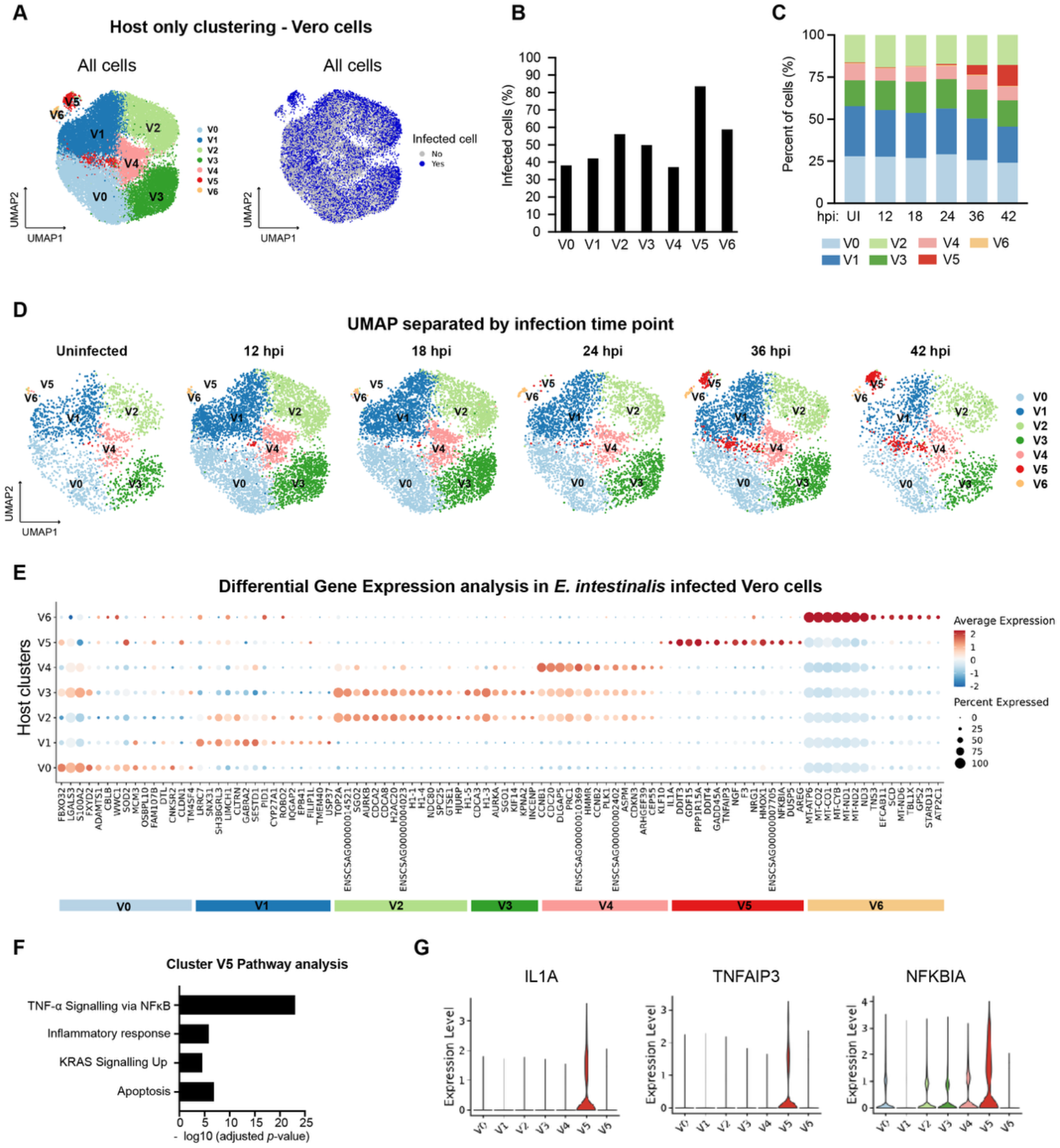
Analysis of host transcriptional response to *E. intestinalis* infection in Vero cells. **A.** UMAP plot of host-only transcripts for all cells (left) and colored by uninfected cells and infected cells (right) across all time points. **B**. Quantification of the percentage of infected cells in each host cluster (n=24,543 cells). **C**. Quantification of the proportion of cells per cluster at 12, 18, 24, 36 and 42 hpi along with uninfected (UI) control cells. **D**. UMAP plot of host-only transcripts separated by infection time point. **E**. Dotplot of the top 15 differentially expressed genes in each host cluster. The X-axis shows the genes and the Y-axis shows the host clusters. The color represents the average gene expression level across the cells in each host cluster in which it is expressed and the size indicates the percentage of cells in each host cluster expressing that gene. **F**. Bar plots showing −log_10_ adjusted *p-*value from gene set enrichment analysis of biological pathways on host cluster V5 using the hallmark gene set in the Human Molecular Signatures Database (MSigDB) [31,32]. **G**. Violin plots showing the expression levels for the top differentially expressed genes in host cluster V5.

## Discussion

Host responses to microsporidia infection vary greatly depending on the microsporidian species, with a general cellular shutdown in response to *T. hominis* infection [14], an upregulation in antimicrobial peptides following *N. bombycis* infection of silkworms [28] and a role for ubiquitin mediated defense in response to *N. parisii* infection in *C. elegans* [10]. In macrophages [15], Caco-2 cells, and Vero cells, *E. intestinalis* infection remains largely undetected during the early stages of parasite development, leading to very little change at the transcriptional level in the majority of cells. However, at least one cluster was induced upon infection in both cell lines: cluster C5 in Caco-2 cells and cluster V5 in Vero cells (Figure 4C-D and 5C-D). Cells in these clusters had a high rate of infection, appeared at later infection time points (36 hpi onwards) and were transcriptionally distinct from other clusters, with the induction of genes involved in TNF-α signalling via NF-κB. This, together with the presence of parasites primarily in the later stages of development, suggest that cells in these clusters were detecting infection and mounting an inflammatory response. A similar transcriptional signature was also observed in *E. intestinalis* infected macrophages (referred to as cluster H5 in [15]) and might represent a shared host response pathway for the detection of later stage parasites in order to limit infection. A second cluster C2 was also observed in Caco-2 cells at later infection time points, though cells in this cluster were not as heavily infected (~40%) and only 23% of parasites in this cluster were in the latest stage of development (P5). However, these cells did display a transcriptional signature similar to cells in cluster C5, suggesting that they might be responding to the same signalling events. In macrophages, a second induced cluster containing a mix of infected and uninfected cells was also observed, which seemed to be responding to interferon signalling as well as TNF-α [15]. We did not observe the induction of any other host clusters in response to *E. intestinalis* infection in Vero cells, which might be due to the inability of Vero cells to produce type I interferons.

The transcriptional atlas of *E. intestinalis* development has previously been defined from scRNA-seq data of infected macrophages [15], which identified 5 parasite developmental stages (P0-P4) characterized by distinct molecular markers. Our analyses of the *E. intestinalis* lifecycle in Caco-2 and Vero cells revealed a similar developmental trajectory. We identified cluster P3 that likely represents proliferative cells undergoing binary fission. This state correlates with an increase in parasite burden as indicated by the 3.5 fold increase in parasite transcripts, suggesting an expansion of replicative stage parasites, which is supported by our fixed cell data (Figure 1A). Cluster P4 marks a transition towards spore development and the formation of sporonts, with the upregulation of genes necessary for spore wall development including spore wall protein 1 (found in the expospore), endospore protein 1 (found in the endospore) and a chitooligosaccharide deacetylase-like protein that is important for the deacetylation of chitin [29] (a component of the spore wall) and is implicated in spore wall formation in *E. cuniculi* [30]. The last parasite developmental stage observed in our dataset, P5, was characterized by an upregulation in genes encoding the polar tube

## Acknowledgements

We thank Kacie McCarty, Olivia Choi, Monica Perumattam, Nicolas Coudray and Pattana Jaroenlak for feedback on the manuscript and all members of the Bhabha+Ekiert labs for helpful discussions. We gratefully acknowledge the following funding source, Johns Hopkins Catalyst Award, awarded to Gira Bhabha. We thank the Integrated Imaging Center at JHU for providing access to their microscopes. Library preparation and 10X scRNA-seq was performed in the Single Cell and Functional Genomics facility of the Integrated Genomics Core of the Johns Hopkins School of Medicine, RRID:SCR_018669. Data analysis was carried out at the Advanced Research Computing at Hopkins (ARCH) core facility (rockfish.jhu.edu), which is supported by the National Science Foundation (NSF) grant number OAC1920103.

## Data Availability

All sequencing data is deposited in NCBI’s Gene Expression Omnibus (GSE347271). Light microscopy data will be made available on Zenodo. A large language model (claude sonnet 4.5, hopGPT) was used for code scaffolding where indicated in the methods section. All code to perform this analysis and generate figures is available on GitHub (https://github.com/nantao1/scRNAseq_Caco2_Vero_E_intestinalis). proteins (PTP1, PTP2, PTP3, PTP4 and PTP6), along with proteins containing Ricin-B domains. This cluster likely represents sporoblasts, a developmental stage during which the spore wall is present and the polar tube is being built.

Taken together, our data suggest conserved core host inflammatory signalling response pathways that are induced in response to *E. intestinalis* infection in multiple cell lines. The parasite development trajectory is similar across the different cell lines tested, with small differences in replicative burden and parasite gene expression being observed. The scRNA-seq datasets generated are a useful resource for correlating future lines of experimentation focused on *E. intestinalis* infection.

## Methods

### Maintenance of mammalian cell lines

Vero cells (ATCC CCL-81) were grown in Dulbecco’s Modified Eagle Medium (DMEM, high glucose, ThermoFisher Scientific 11965092) supplemented with 10% heat-inactivated fetal bovine serum (FBS) (VWR Life Science; 89510-188) and 1X MEM non-essential amino acids solution (Gibco− 11140050). Caco-2 cells (ATCC HTB-37^−^) were grown in DMEM, high glucose growth medium supplemented with 20% FBS and 1X MEM non-essential amino acids solution. Cells were grown at 37 °C in a humidified atmosphere with 5% CO2. Cell lines are tested each month using the Venor®GeM Classic Mycoplasma Detection Kit (minerva biolabs 11-1100G) and found negative.

### Propagation of E. intestinalis spores

*E. intestinalis* (ATCC 50506) spores were propagated in Vero cells. Vero cells were cultured in DMEM, high glucose with 10% heat-inactivated FBS and 1X MEM non-essential amino acids solution at 37 °C and with 5% CO_2_. Vero cells were seeded at 2.1E6 in a 75 cm^2^ flask. After two days the media was switched to DMEM, high glucose supplemented with 3% FBS and 1X MEM non-essential amino acids solution and *E. intestinalis* spores were added. The media was changed every 2 days for 10 days before purification of mature *E. intestinalis* spores.

### Purification of E. intestinalis spores

Purification was carried out using a detailed protocol we have previously published [33]. Briefly, infected cells were detached from tissue culture flasks using a cell scraper and placed into a 50 mL conical tube, followed by centrifugation at 2000 ×*g* for 10 min at 25 °C. Cells were resuspended in sterile DPBS (ThermoFisher Scientific 14190250) with 0.1% SDS, vortexed and incubated on ice for 10 min. The released spores were then purified using a continuous Percoll gradient. Equal volumes of spore suspension and 100% Percoll were added to a 15 mL conical tube, vortexed and then centrifuged at 2000 ×*g* for 30 min at room temperature. The purified spore pellets were washed three times with 10 mL sterile DPBS and further purified in a discontinuous Percoll gradient. Briefly, spore pellets were resuspended in 2 mL of sterile DPBS and layered onto a 10 mL percoll gradient (2.5 mL 100% Percoll, 2.5 mL 75% Percoll, 2.5 mL 50% Percoll, 2.5 mL 25% Percoll) in a 15 mL conical tube, and centrifuged at 7000 ×*g* for 30 min at RT in a Bioflex HC rotor. Spores that separated into the pellet (100% Percoll) were carefully removed and washed twice with 10 mL of sterile DPBS, centrifuging at 2000 ×*g* for 5 min at RT in between washes. Purified spore pellets were stored in sterile DPBS at 4 °C for further analyses.

### E. intestinalis infection experiments

To collect data for each infection experiment two parallel sets of cultures were prepared for *E. intestinalis*-infected Caco-2 cells or *E. intestinalis*-infected Vero cells. One set of cultures was processed for scRNA-seq, and the second set of cultures was prepared for fluorescence microscopy to assess the infection levels at each timepoint. For Caco-2 cells, 3.3 x 10^5^ cells were seeded into a 6-well plate that was empty (scRNA-seq) or contained five 12 mm circular glass coverslips per well (microscopy). Caco-2 cells were allowed to rest for 48 h at 37 °C with 5% CO2 in DMEM, high glucose with 20% FBS, 1X MEM non-essential amino acids. For Vero cells, 1 x 10^5^ cells were seeded into a 6-well plate that was empty (scRNA-seq) or contained five 12 mm circular glass coverslips per well (microscopy). Vero cells were allowed to rest for 24 h at 37 °C with 5% CO2 in DMEM, high glucose with 10% FBS, 1X MEM non-essential amino acids. Prior to infection with *E. intestinalis*, old media was removed and fresh media was added to each well. To infect cells, *E. intestinalis* spores were added at an MOI 30 for 42 h, 36 h, 24 h, 18 h, 12 h after which they were harvested for scRNA-seq analyses or microscopy assessment of infection.

### RNA FISH and Fluorescence microscopy

To assess *E. intestinalis* infection and replication in Vero and Caco-2 cells, media was removed from each well and cells on coverslips were washed 1x in PBS (148.8 mM NaCl, 9.5 mM Na_2_HPO_4_.7H_2_O, 2.6mM KCl, 1.4mM KH_2_PO_4_) and fixed in 4% PFA in PBS-T (0.1% Tween-20) at 37 °C for 15 min. Coverslips were washed 2x with 2 mL PBS-T and 1x with 500 µL hybridization buffer (900 mM NaCl, 20 mM Tris HCl pH 7.5, 0.01% SDS). Coverslips were transferred face up to a hybridization chamber and 50 μL FISH staining solution (125 nM FISH probe in hybridization buffer) was added per coverslip and incubated for 18 h at 37 °C in the dark. An *E. intestinalis* 16S rRNA-specific FISH probe conjugated to Quasar 590 (designed using the Stellaris RNA FISH Probe Designer Biosearch Technologies, Inc., Petaluma, CA) was used [33]. Following incubation, coverslips were washed 2x with 500 µL of wash buffer (50 mL hybridization buffer + 5 mM EDTA) for 30 min at 37 °C to remove excess FISH probe. Cells were washed 1x with PBS-T and then incubated with a staining solution containing three dyes - calcofluor white (18909-100ML-F), DRAQ5 (NBP2-81125-50ul), and Phalloidin 488 (Cayman Chemical Company 20549). Final conditions for the dyes were: 2.5 µg/mL calcofluor white, 5uM DRAQ5, 1X Phalloidin 488 in PBS-T). Samples were incubated for 30 min at RT. Coverslips were washed 1x with PBS-T and mounted onto slides with Prolong Diamond antifade (ThermoFisher Scientific; P36965) and sealed. Samples were imaged on a Nikon W1 spinning disc confocal microscope with a Nikon Plan Apo 60x 1.42 Oil objective and a Hamamatsu Orca Fusion-BT camera. Z-stacks were acquired with 0.2 μm spacing. Maximum intensity projection images were created in the Nikon Elements software. Pseudo coloring, cropping and analysis were performed in Fiji [34] and images were assembled in Illustrator (Adobe, San Jose, CA). 1,917 cells were scored for the Caco-2 infected-cell dataset and 1,889 cells were scored for the Vero infected-cell dataset. Cells were scored as infected if they were positive for RNA FISH signal only (early parasite developmental stages) or RNA FISH signal and Calcofluor white signal (mid-late parasite developmental stages).

### Generating single-cell suspensions and cell hashing

To generate single cells for scRNA-seq, infected Caco-2 or Vero cells were detached from the 6-well plate by trypsinization. Briefly, media was removed from the well, cells were washed with 2 mL of sterile DPBS and incubated with 200 uL of 0.25% trypsin, 2.21 mM EDTA solution (Corning 25-053-CI) at 37 °C with 5% CO_2_ for 5 min. 800 uL DMEM, high glucose media supplemented with 1X MEM non-essential amino acids and either 20% FBS (Caco-2 cells) or 10% FBS (Vero cells) was added to stop trypsinization. Cells were resuspended and further dissociated by pipetting 4-5x. Cell suspension was transferred to a 1.5 mL tube and centrifuged at 200 × g for 5 min at 4 °C.

To combine cells from different infection timepoints and uninfected cells for scRNA-seq library preparation, the cells from each infection timepoint were incubated with different hashing antibodies [35]. Cells were resuspended in 100 μl of a cell staining buffer (2% BSA and 0.01% Tween 20 in 1X PBS). 10 μl of Fc blocking reagent (BioLegend, USA) was added to each reaction, and incubated for 10 min at 4 °C. 0.5 µg of TotalSeq B anti-human hashtag antibodies TotalSeq-B5-10 (BioLegend, USA) was added into the cell suspensions collected from each infection time point and incubated for 25 min at 4 °C. The cells were washed three times with 1 ml of staining buffer, centrifuging at 1000 *x g* for 5 min at 4 °C in between washes. Cell pellets were resuspended in 150 μl of PBS supplemented with 10% FBS. Cell density and viability were counted following staining with AOPI on the Cellometer Ascend Automated Cell Counter. Cell viability for each infection time point ranged from 88-95% for Caco-2 cells and 94-95% for Vero cells.

### Library preparation and 10X scRNA-seq

After antibody hashing, cells were pooled as follows: 5000 cell target for 12 hpi and 18 hpi, 3333 cell target for 24 hpi and 36 hpi, 2000 cells for 42 hpi, and 1500 cells for the uninfected sample, for a 20,000 cell target per lane. Libraries were generated according to the manufacturer’s protocol (10x Genomics, USA) and the antibody hash library was amplified with 8 cycles. Libraries were pooled at a 5:1 ratio for GEX:HASH, respectively and sequenced on half a lane of a NovaSeq X 10B flow cell (Illumina, USA). Library preparation and sequencing were carried out at the Single Cell & Functional Genomics at the Integrated Genomics Center (IGC), at Johns Hopkins.

### Processing of the raw sequencing reads

A combined reference genome containing both the human genome (GRch38; GCF_000001405.40; contains nuclear and mitochondrial genomes) and the *E. intestinalis* genome (ATCC 50506; GCF_000146465.1) or the African Green Monkey genome (GCF_000409795.2) and the *E. intestinalis* genome was generated using Cell Ranger software version 9.0.1 with the Cell Ranger *mkref* function (10X Genomics, USA). No quality control was performed prior to Cell Ranger. Raw sequencing reads were mapped to the combined reference genome and the gene expression matrices were generated using the Cell Ranger *count* function with default parameters. Cell Ranger *count* quantifies the amount of genes detected in each cell (nFeature) and the total number of the RNA molecules in each cell (nCount), which was evaluated from unique molecular identifiers (UMIs).

## scRNA-seq data processing

### Initial data processing

The gene expression matrices from Cell Ranger *count* were transferred and processed using Seurat version 5.3.0. Percentage mitochondria in each cell was calculated using the PercentageFeatureSet, pattern = = “MT-”. To identify which cells were infected with *E. intestinalis*, the percentage of *E. intestinalis* transcripts detected in each cell were calculated (PercentageFeatureSet, pattern = “^EI50506-”). We classified cells that have >0.2% of the microsporidian transcripts as “infected”. Varying this cutoff between 0.2% and 80% microsporidian transcripts drastically changed the number of cells being scored as infected in the infected samples but did not drastically increase the number of cells being scored as infected in the uninfected control samples (Supplementary Figure 1C&1E). The number of infected cells found in each dataset are shown in Supplementary Figure 1D, 1F and 2C. A small portion of the control cells are classified as “infected”, even though no parasites were added to these samples; similar results were observed in previous scRNA-seq of *E. intestinalis*-infected macrophages. In light of the fact that we pooled our uninfected cells together with all the infected cell timepoints just before loading into the Chromium Controller, we hypothesize that these are actually rare doublet events due to co-encapsulation of an uninfected control cell (with “control” cell hashing barcode), and an infected cell/debris without a detectable cell hashing barcode, which our pipeline cannot readily remove. Cells from different infection timepoints were demultiplexed based on their hashing antibodies, using a HTODemux function. Doublet cells, which contained more than one type of hashing antibody, and negative cells, which lacked any hashing antibody were removed from all of the datasets. To further filter low quality cells, nFeature, nCount, and percentage of mitochondrial genes (percent.mt) were used. In the Caco-2 infected cell dataset, the criteria were nFeature: 400-6500 genes, nCount: 1000-30,000 UMIs, and percent.mt <20. For the Vero infected-cell dataset, the criteria were nFeature: 400–4500 genes, nCount: 1000-16,000 UMIs, and percent.mt <20. The number of cells obtained from each cell line after filtering are shown in Supplementary Figure 1A and B.

### scRNA-seq data processing of the human and Chlorocebus sabaeus transcripts

The human and Chlorocebus sabaeus transcripts were separated from *E. intestinalis* transcripts by a subset function prior to normalization. The datasets were then normalized using a log normalization method (NormalizeData, normalization.method = “LogNormalize”,scale.factor = 10,000) and variable features were identified(FindVariableFeatures,selection.method = “vst”,nfeatures = 3000). Then, we ran RunPCA, PC1 to PC42 (Caco-2 infected cell dataset) and PC1 to PC43 (Vero infected cell dataset) were included to perform dimensionality reduction using a Uniform Manifold Approximation and Projection (UMAP) method (RunUMAP, dims = 1:42 for Caco-2 infected cell dataset and 1:43 for Vero infected cell dataset). For cell clustering, FindNeighbors (dims = 1:42 or 1:43) and FindClusters (resolution = 0.25 for Caco-2 infected dataset and 0.2 for Vero infected cell dataset) were carried out. DimPlot and FeaturePlot function were used to generate UMAP plots in Figures 1,4,5 and Supplementary Figure 4.

### scRNA-seq data processing of the E. intestinalis transcripts

*E. intestinalis* transcripts were separated from the human or Chlorocebus sabaeus transcripts by a subset function prior to normalization. Only infected cells identified from the initial scRNA-seq data processing were used in this analysis. RunPCA was performed followed by RunUMAP PC1 to PC43 (RunUMAP, dims = 1:43) for both datasets. To cluster the infected cells, FindNeighbors (dims = 1:43) and FindClusters (resolution = 0.4) were performed. DimPlot and FeaturePlot function were used to generate plots in Figures 2-3 and Supplementary Figure 2.

### Marker gene detection and differential gene expression analyses

To identify signature genes for each cell cluster, FindAllMarkers function was performed using the default parameters and the min.cpt of 0.25 (FindAllMarkers, min.pct = 0.25). Wilcoxon Rank Sum test was used to test the differentially expressed genes between clusters. Average Log_2_ Fold change (avg_log2FC) was also used to rank highly expressed genes in each cluster. DotPlot was carried out to generate Figures 2C, 3C, 4E, 5E, S1G and S1H. VlnPlot was used to visualize the gene expression levels in Figures 2D, 3D, 4G, and 5G.

### Gene set enrichment analysis

To better understand which pathways highly expressed genes in cluster C5, C2, C3 and V5 are involved in, pathway enrichment analysis was performed using KEGG [36–38] and fGSEA pathway enrichment tools in R. Briefly, highly expressed genes that are upregulated and downregulated, with a fold change >1.5 were selected and subjected to KEGG pathway (enrichKEGG) and fGSEA (fgsea(pathways = hallmark_list)) analysis in R. Figures 4F, 5F, and S4C were obtained using the ‘hallmark gene sets’ in the Human Molecular Signatures Database (MSigDB) [31,32]. The terms were ranked according to their *p*-values. This code was generated with the help of claude sonnet 4.5 made available through hopGPT (JHU), verified and modified by the authors.

### Comparing top differentially expressed genes during parasite development across macrophage, Caco-2 and Vero cells

To identify cell-type specific differences in gene expression across parasite developmental stages (P0-P5 for Caco-2 and Vero cells; P0-P4 for macrophages), we identified the top 50 differentially expressed genes at each stage for each cell line. Using tidyverse (dplyr, tidyr) packages in R, pairwise comparisons e.g. Caco-2 vs Macrophage P0, were generated using the full join function to retain all the genes from each dataset. The left join function was then used to incorporate average expression data (pseudobulking Seurat - AverageExpression function). Log2 average expression values were generated and the data pivoted into a wide format with individual genes as rows and cell lines as columns. Linear regression models were fitted to the data to assess how well correlated gene expression was across the different cell lines. Finally, ggplot2 was used to generate scatter plots. This code was generated with the help of claude sonnet 4.5 made available through hopGPT (JHU), verified and modified by the authors.

## References

1. Han B, Pan G, Weiss LM. 2021 Microsporidiosis in humans. Clin. Microbiol. Rev. 34, e0001020. (doi:10.1128/CMR.00010-20)

2. Orenstein JM. 1996 Intestinal Microsporidiosis. Advances in Anatomic Pathology. 3, 46–58. (doi:10.1097/00125480-199601000-00050)

3. Kechaou R et al. 2025 Microsporidiosis in patients with autoimmune diseases undergoing monoclonal antibody associated therapy. Mycopathologia 190, 12. (doi:10.1007/s11046-024-00918-2)

4. Desportes I, Le Charpentier Y, Galian A, Bernard F, Cochand-Priollet B, Lavergne A, Ravisse P, Modigliani R. 1985 Occurrence of a new microsporidan: Enterocytozoon bieneusi n.g., n. sp., in the enterocytes of a human patient with AIDS. J. Protozool. 32, 250–254. (doi:10.1111/j.1550-7408.1985.tb03046.x)

5. Sandfort J, Hannemann A, Gelderblom H, Stark K, Owen RL, Ruf B. 1994 Enterocytozoon bieneusi Infection in an Immunocompetent Patient Who Had Acute Diarrhea and Who Was Not Infected with the Human Immunodeficiency Virus. Clinical Infectious Diseases. 19, 514–516. (doi:10.1093/clinids/19.3.514)

6. Han B et al. 2017 The role of microsporidian polar tube protein 4 (PTP4) in host cell infection. PLoS Pathog. 13, e1006341. (doi:10.1371/journal.ppat.1006341)

7. Cali A, Kotler DP, Orenstein JM. 1993 Septata intestinalis N. G., N. Sp., an intestinal microsporidian associated with chronic diarrhea and dissemination in AIDS patients. J. Eukaryot. Microbiol. 40, 101–112. (doi:10.1111/j.1550-7408.1993.tb04889.x)

8. Canning EU, Field AS, Hing MC, Marriott DJ. 1994 Further observations on the ultrastructure of Septata intestinalis Cali, Kotler and Orenstein, 1993. Eur. J. Protistol. 30, 414–422. (doi:10.1016/s0932-4739(11)80216-6)

9. Antao NV et al. 2023 3D reconstructions of parasite development and the intracellular niche of the microsporidian pathogen Encephalitozoon intestinalis. Nat Commun 14, 7662. (doi:10.1038/s41467-023-43215-0)

10. Bakowski MA, Desjardins CA, Smelkinson MG, Dunbar TL, Lopez-Moyado IF, Rifkin SA, Cuomo CA, Troemel ER. 2014 Ubiquitin-mediated response to microsporidia and virus infection in C. elegans. PLoS Pathog 10, e1004200. (doi:10.1371/journal.ppat.1004200)

11. Grisdale CJ, Bowers LC, Didier ES, Fast NM. 2013 Transcriptome analysis of the parasite Encephalitozoon cuniculi: an in-depth examination of pre-mRNA splicing in a reduced eukaryote. BMC Genomics 14, 207. (doi:10.1186/1471-2164-14-207)

12. Wan YC, Troemel ER, Reinke AW. 2022 Conservation of Nematocida microsporidia gene expression and host response in Caenorhabditis nematodes. PLoS One 17, e0279103. (doi:10.1371/journal.pone.0279103)

13. Flores J, Takvorian PM, Weiss LM, Cali A, Gao N. 2021 Human microsporidian pathogen impinges on enterocyte membrane trafficking and signaling. J Cell Sci 134. (doi:10.1242/jcs.253757)

14. Watson AK, Williams TA, Williams BAP, Moore KA, Hirt RP, Embley TM. 2015 Transcriptomic profiling of host-parasite interactions in the microsporidian Trachipleistophora hominis. BMC Genomics 16, 983. (doi:10.1186/s12864-015-1989-z)

15. Jaroenlak P et al. 2025 scRNA-seq uncovers the transcriptional dynamics of Encephalitozoon intestinalis parasites in human macrophages. Nat Commun 16, 3269. (doi:10.1038/s41467-025-57837-z)

16. Lea T. 2015 Caco-2 cell line.

17. Taddeo B, Esclatine A, Zhang W, Roizman B. 2003 The stress-inducible immediate-early responsive gene IEX-1 is activated in cells infected with herpes simplex virus 1, but several viral mechanisms, including 3’ degradation of its RNA, preclude expression of the gene. J Virol 77, 6178–6187. (doi:10.1128/jvi.77.11.6178-6187.2003)

18. Arlt A, Schäfer H. 2011 Role of the immediate early response 3 (IER3) gene in cellular stress response, inflammation and tumorigenesis. Eur J Cell Biol 90, 545–552. (doi:10.1016/j.ejcb.2010.10.002)

19. Vereecke L, Beyaert R, van Loo G. 2009 The ubiquitin-editing enzyme A20 (TNFAIP3) is a central regulator of immunopathology. Trends Immunol 30, 383–391. (doi:10.1016/j.it.2009.05.007)

20. Ohmae T, Hirata Y, Maeda S, Shibata W, Yanai A, Ogura K, Yoshida H, Kawabe T, Omata M. 2005 Helicobacter pylori activates NF-kappaB via the alternative pathway in B lymphocytes. J Immunol 175, 7162–7169. (doi:10.4049/jimmunol.175.11.7162)

21. Ge J, Xu H, Li T, Zhou Y, Zhang Z, Li S, Liu L, Shao F. 2009 A Legionella type IV effector activates the NF-kappaB pathway by phosphorylating the IkappaB family of inhibitors. Proc Natl Acad Sci U S A 106, 13725–13730. (doi:10.1073/pnas.0907200106)

22. Rückle A, Haasbach E, Julkunen I, Planz O, Ehrhardt C, Ludwig S. 2012 The NS1 protein of influenza A virus blocks RIG-I-mediated activation of the noncanonical NF-κB pathway and p52/RelB-dependent gene expression in lung epithelial cells. J Virol 86, 10211–10217. (doi:10.1128/JVI.00323-12)

23. Onose A et al. 2006 An inhibitory effect of A20 on NF-kappaB activation in airway epithelium upon influenza virus infection. Eur J Pharmacol 541, 198–204. (doi:10.1016/j.ejphar.2006.03.073)

24. Hitotsumatsu O et al. 2008 The ubiquitin-editing enzyme A20 restricts nucleotide-binding oligomerization domain containing 2-triggered signals. Immunity 28, 381–390. (doi:10.1016/j.immuni.2008.02.002)

25. Scanlon M, Shaw AP, Zhou CJ, Visvesvara GS, Leitch GJ. 2000 Infection by microsporidia disrupts the host cell cycle. J. Eukaryot. Microbiol. 47, 525–531. (doi:10.1111/j.1550-7408.2000.tb00085.x)

26. Dube PH, Revell PA, Chaplin DD, Lorenz RG, Miller VL. 2001 A role for IL-1 alpha in inducing pathologic inflammation during bacterial infection. Proc Natl Acad Sci U S A 98, 10880–10885. (doi:10.1073/pnas.191214498)

27. Di Paolo NC, Shayakhmetov DM. 2016 Interleukin 1α and the inflammatory process. Nat Immunol 17, 906–913. (doi:10.1038/ni.3503)

28. Ma Z et al. 2013 Genome-wide transcriptional response of silkworm (Bombyx mori) to infection by the microsporidian Nosema bombycis. PLoS One 8, e84137. (doi:10.1371/journal.pone.0084137)

29. Pochanavanich P, Suntornsuk W. 2002 Fungal chitosan production and its characterization. Lett. Appl. Microbiol. 35, 17–21. (doi:10.1046/j.1472-765x.2002.01118.x)

30. Brosson D, Kuhn L, Prensier G, Vivarès CP, Texier C. 2005 The putative chitin deacetylase of Encephalitozoon cuniculi: a surface protein implicated in microsporidian spore-wall formation. FEMS Microbiol. Lett. 247, 81–90. (doi:10.1016/j.femsle.2005.04.031)

31. Subramanian A et al. 2005 Gene set enrichment analysis: a knowledge-based approach for interpreting genome-wide expression profiles. Proc. Natl. Acad. Sci. U. S. A. 102, 15545–15550. (doi:10.1073/pnas.0506580102)

32. Liberzon A, Birger C, Thorvaldsdóttir H, Ghandi M, Mesirov JP, Tamayo P. 2015 The Molecular Signatures Database (MSigDB) hallmark gene set collection. Cell Syst. 1, 417–425. (doi:10.1016/j.cels.2015.12.004)

33. Noelle V. Antao, Mahrukh Usmani, Frederick Rubino, Harshita Ramchandani, Pattana Jaroenlak, Kacie L. McCarty, Damian C. Ekiert, Gira Bhabha. 07 2024 Purification of germination-competent E. intestinalis spores. protocols. io (doi:10.17504/protocols.io.3byl495wogo5/v1)

34. Schindelin J et al. 2012 Fiji: an open-source platform for biological-image analysis. Nat. Methods 9, 676–682. (doi:10.1038/nmeth.2019)

35. Stoeckius M, Zheng S, Houck-Loomis B, Hao S, Yeung BZ, Mauck WM 3rd, Smibert P, Satija R. 2018 Cell Hashing with barcoded antibodies enables multiplexing and doublet detection for single cell genomics. Genome Biol. 19, 224. (doi:10.1186/s13059-018-1603-1)

36. Kanehisa M, Goto S. 2000 KEGG: kyoto encyclopedia of genes and genomes. Nucleic Acids Res. 28, 27–30. (doi:10.1093/nar/28.1.27)

37. Kanehisa M. 2019 Toward understanding the origin and evolution of cellular organisms. Protein Sci. 28, 1947–1951. (doi:10.1002/pro.3715)

38. Kanehisa M, Furumichi M, Sato Y, Matsuura Y, Ishiguro-Watanabe M. 2025 KEGG: biological systems database as a model of the real world. Nucleic Acids Res. 53, D672–D677. (doi:10.1093/nar/gkae909)

