## Supplementary Figures for "Single-cell transcriptomics of *Encephalitozoon intestinalis* infection in Caco-2 and Vero cells"

### Supplementary Figure 1

**A**

Cell counts Caco-2 infected cell dataset

| Host Clusters | 12 hpi | 18 hpi | 24 hpi | 36 hpi | 42 hpi | Uninfected | Total |
|---|---|---|---|---|---|---|---|
| C0 | 3116 | 2682 | 1445 | 136 | 115 | 926 | 8420 |
| C1 | 2111 | 1978 | 1366 | 333 | 170 | 731 | 6689 |
| C2 | 342 | 294 | 162 | 2816 | 1444 | 152 | 5210 |
| C3 | 351 | 351 | 226 | 195 | 20 | 195 | 1338 |
| C4 | 185 | 162 | 84 | 51 | 24 | 50 | 556 |
| C5 | 0 | 0 | 0 | 174 | 129 | 0 | 303 |
| C6 | 28 | 43 | 51 | 45 | 25 | 13 | 205 |
| C7 | 6 | 6 | 5 | 6 | 2 | 2 | 27 |
| | | | | | | | 22748 |

**B**

Cell counts Vero infected cell dataset

| Host Clusters | 12 hpi | 18 hpi | 24 hpi | 36 hpi | 42 hpi | Uninfected | Total |
|---|---|---|---|---|---|---|---|
| V0 | 1607 | 1647 | 1115 | 1108 | 575 | 591 | 6643 |
| V1 | 1602 | 1636 | 1043 | 1070 | 513 | 625 | 6489 |
| V2 | 1111 | 1128 | 659 | 771 | 424 | 344 | 4437 |
| V3 | 1007 | 1136 | 668 | 743 | 374 | 326 | 4254 |
| V4 | 417 | 527 | 297 | 362 | 194 | 206 | 2003 |
| V5 | 21 | 12 | 25 | 246 | 298 | 6 | 608 |
| V6 | 18 | 24 | 24 | 23 | 10 | 10 | 109 |
| | | | | | | | 24543 |

**C**

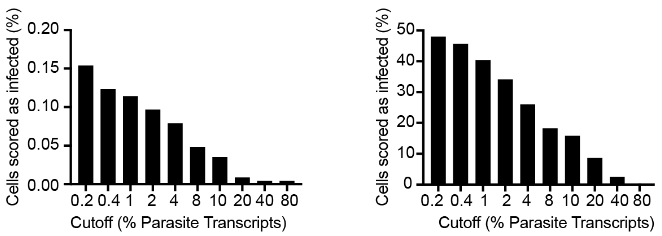


**D**

Caco-2 cells

| Samples | 2% Microsporidia Transcript Cutoff | 0.2% Microsporidia Transcript Cutoff |
|---|---|---|
| 12 hpi | 1634 | 3260 |
| 18 hpi | 2447 | 3167 |
| 24 hpi | 1588 | 1821 |
| 36 hpi | 1372 | 1701 |
| 42 hpi | 701 | 931 |
| Uninfected | 22 | 35 |
| Total | 7764 | 10915 |

**E**

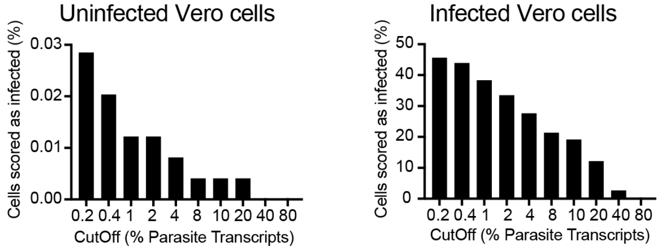


**F**

Vero cells

| Samples | 2% Microsporidia Transcript Cutoff | 0.2% Microsporidia Transcript Cutoff |
|---|---|---|
| 12 hpi | 1310 | 2888 |
| 18 hpi | 2450 | 3284 |
| 24 hpi | 1792 | 2047 |
| 36 hpi | 1716 | 1892 |
| 42 hpi | 946 | 1074 |
| Uninfected | 3 | 7 |
| Total | 8217 | 11192 |

**G**

**Differential Gene Expression analysis of the total transcriptome in *E. intestinalis* infected Caco-2 cells**

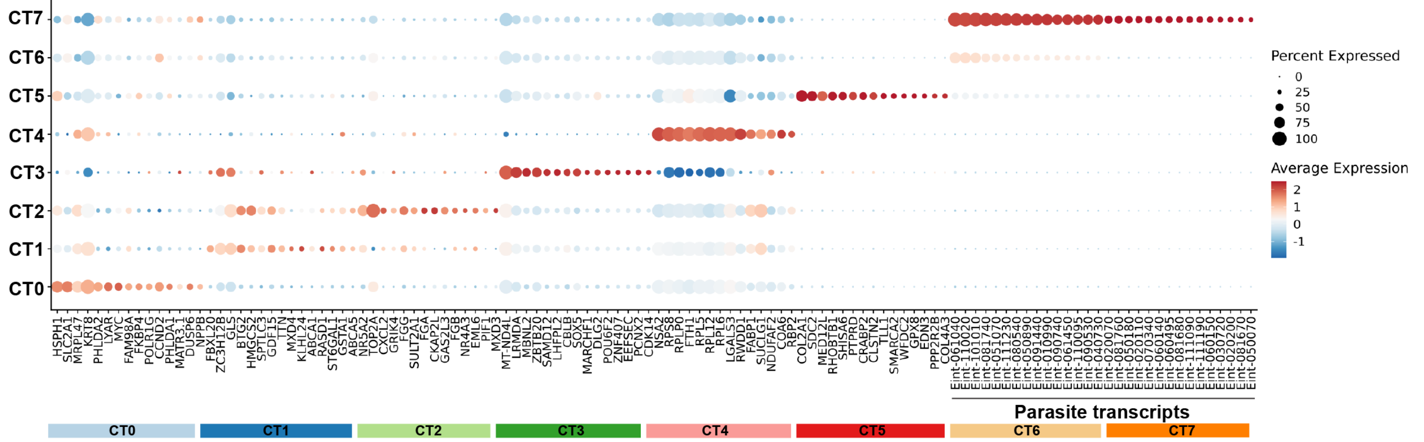


**H**

**Differential Gene Expression analysis of the total transcriptome in *E. intestinalis* infected Vero cells**

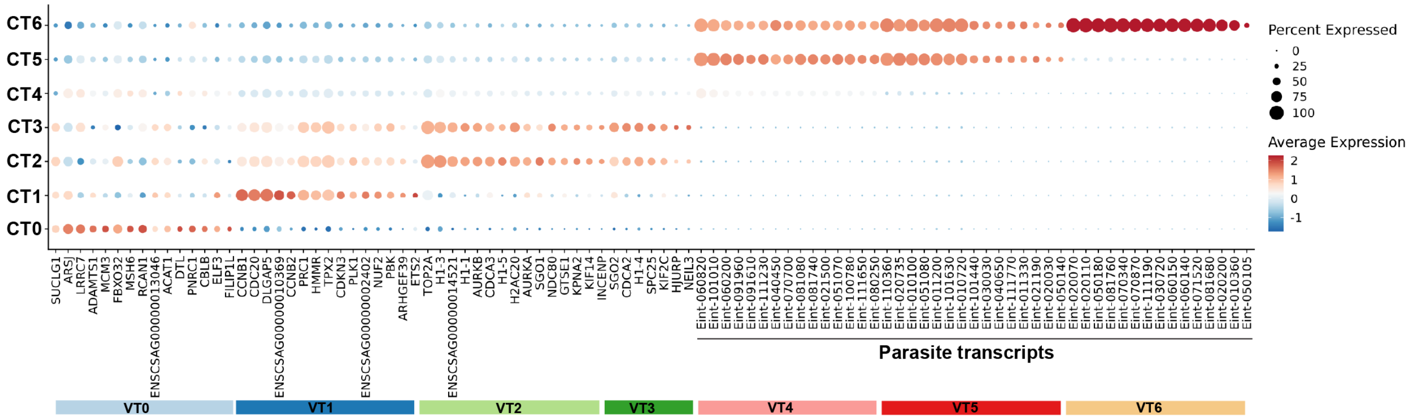

#### Supplementary Figure 2

**A** **Parasite only clustering in Caco-2 cells split by infection time point**

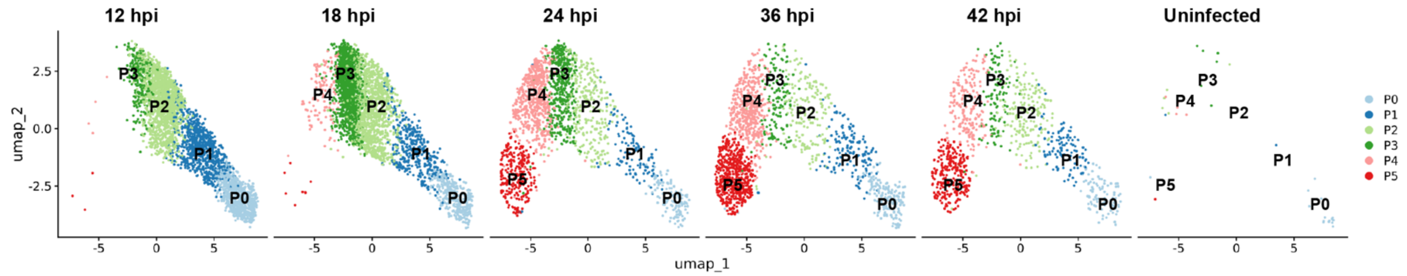


**B** **Caco-2 cells**

| Parasite Cluster | 12 hpi | 18 hpi | 24 hpi | 36 hpi | 42 hpi | Uninfected |
|---|---|---|---|---|---|---|
| P0 | 759 | 354 | 129 | 187 | 93 | 16 |
| P1 | 811 | 377 | 117 | 143 | 105 | 2 |
| P2 | 1442 | 1144 | 276 | 180 | 176 | 5 |
| P3 | 241 | 1162 | 445 | 136 | 102 | 6 |
| P4 | 4 | 122 | 588 | 481 | 197 | 5 |
| P5 | 3 | 8 | 266 | 574 | 258 | 1 |

**C**

| Parasite Cluster | Caco-2 | Vero |
|---|---|---|
| P0 | 1538 | 1838 |
| P1 | 1555 | 1438 |
| P2 | 3223 | 2829 |
| P3 | 2092 | 1906 |
| P4 | 1397 | 1891 |
| P5 | 1110 | 1290 |
| Total | 10915 | 11192 |

**D** **Parasite only clustering in Vero cells split by infection time point**

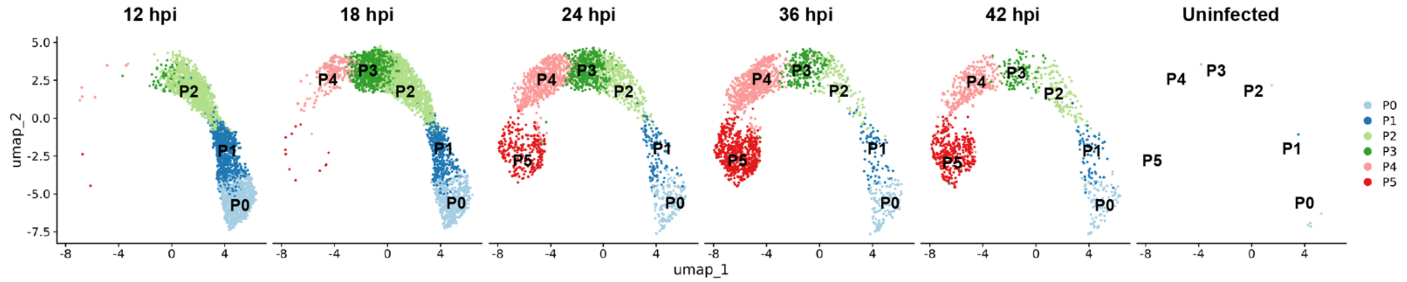


**E** **Vero cells**

| Parasite Cluster | 12 hpi | 18 hpi | 24 hpi | 36 hpi | 42 hpi | Uninfected |
|---|---|---|---|---|---|---|
| P0 | 961 | 486 | 172 | 128 | 87 | 4 |
| P1 | 771 | 409 | 119 | 71 | 67 | 1 |
| P2 | 1070 | 1126 | 342 | 156 | 134 | 1 |
| P3 | 77 | 1005 | 505 | 215 | 104 | 0 |
| P4 | 7 | 246 | 652 | 662 | 323 | 1 |
| P5 | 2 | 12 | 257 | 660 | 359 | 0 |

**Supplementary Figure 3**

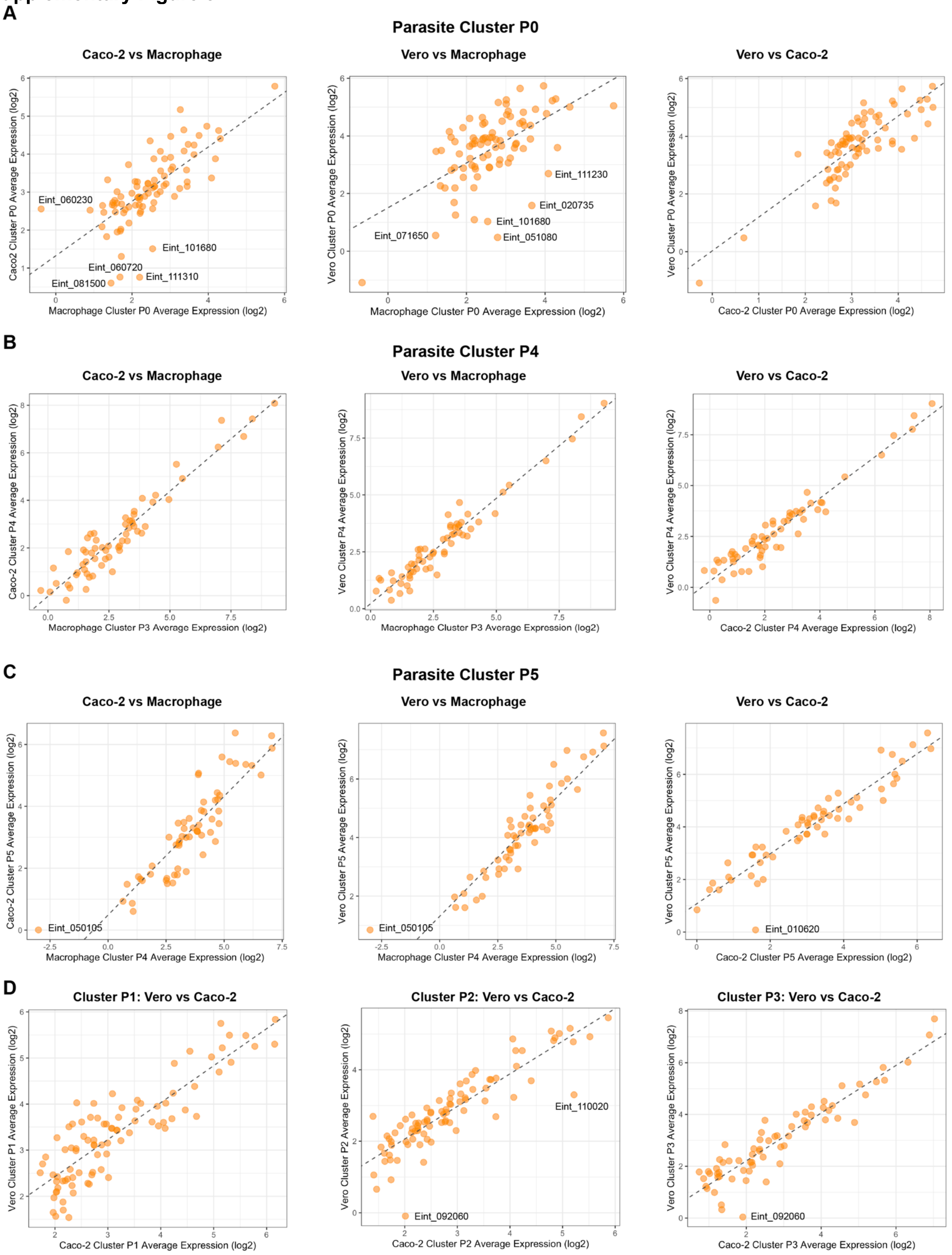

**Supplementary Figure 4**

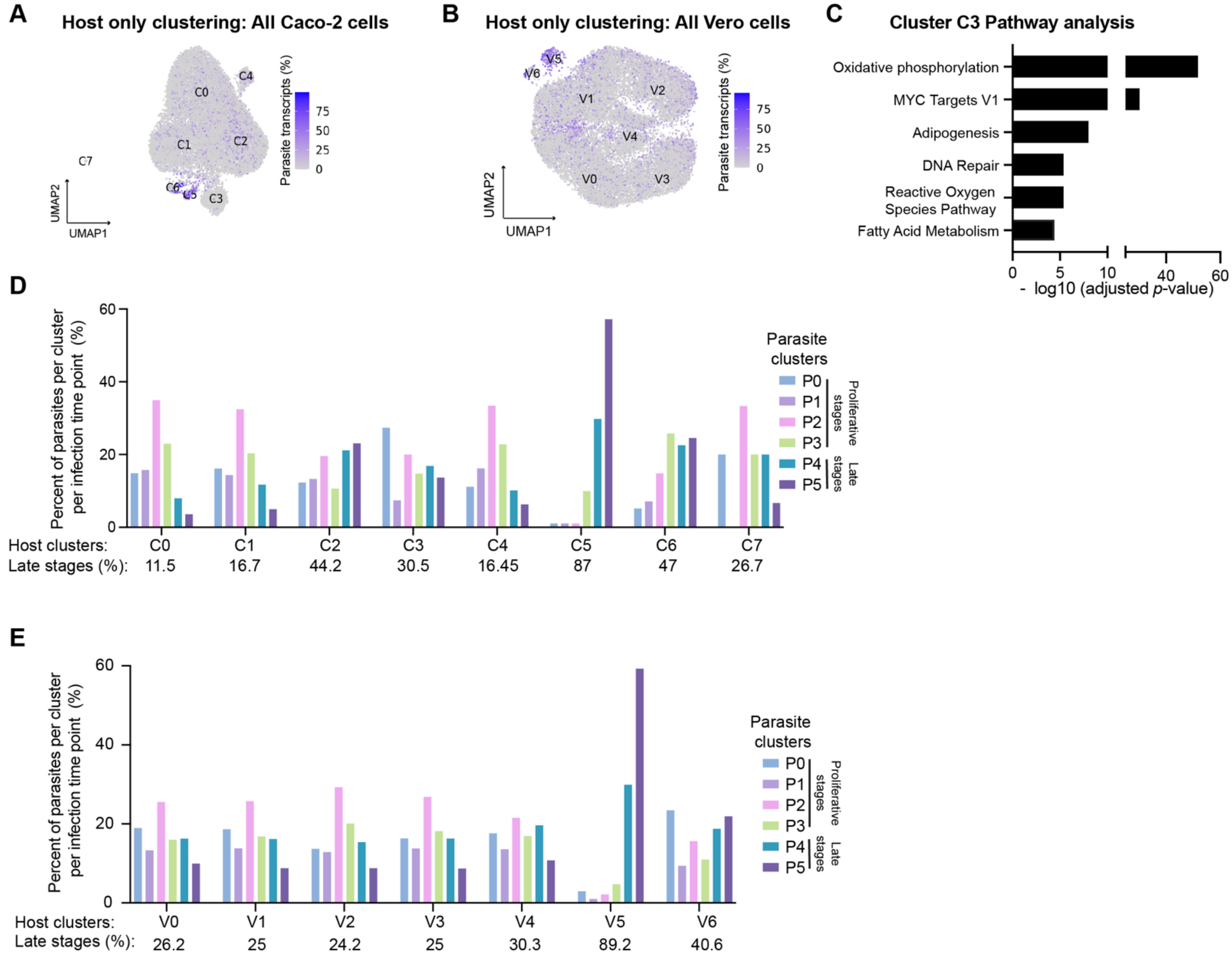

**Supplementary Figure legends**

**Supplementary Figure 1: Quality control and cell filtering for scRNA-seq analysis in Caco-2 and Vero cells. A.** Table of the number of cells at each infection timepoint used in scRNA-seq analysis for Caco-2 infected cell dataset after quality control and cell filtering. **B.**Table of the number of cells at each infection timepoint used in scRNA-seq analysis for the Vero infected cell dataset after quality control and cell filtering. **C.** Graphs plotting the percentage of Caco-2 cells scored as infected with different parasite cutoffs in uninfected control cells and infected cells. **D.** Table of the number of cells at each infection time point with 2% or 0.2% cutoff for the presence of microsporidian transcripts in the Caco-2 infected cell dataset. **E.** Graphs plotting the percentage of Vero cells scored as infected with different parasite cutoffs in uninfected control cells and infected cells. **F.** Table of the number of cells at each infection time point counted as infected based on a 2% or a 0.2% microsporidia transcript cut-off in the Vero infected cell dataset. **G.** Dotplot of the top 15 differentially expressed genes in each Caco-2 host cluster. The X-axis shows the genes and the Y-axis shows the host clusters. The red-blue color range represents the average gene expression level across the cells in each host cluster in which it is expressed and the size indicates the percentage of cells in each host cluster expressing that gene. **H.** Dotplot of the top 15 differentially expressed genes in each Vero host cluster. The X-axis shows the genes and the Y-axis shows the host clusters. The color represents the average gene expression level across the cells in each host cluster in which it is expressed and the size indicates the percentage of cells in each host cluster expressing that gene.

**Supplementary Figure 2: Transcriptional analysis of *E. intestinalis* development in Caco-2 and Vero cells. A.** UMAP plots of parasite-only transcripts from Caco-2 infected cells separated by infection time point. **B.** Table of the number of infected cells (0.2% microsporidia transcript cut-off) found in each parasite developmental cluster separated by infection time point in Caco-2 infected cell dataset. **C.** Table of the number of infected cells found in each parasite developmental cluster in Caco-2 and Vero cells. **D.** UMAP plots of parasite-only transcripts from Vero infected cells separated by infection time point. **E.** Table of the number of infected cells (0.2% microsporidia transcript cut-off) found in each parasite developmental cluster separated by infection time point in Vero infected cell dataset.

**Supplementary Figure 3: Comparison of the top 50 differentially expressed parasite genes at all parasite developmental stages in Caco-2, Vero and Macrophage datasets. A.** Scatter plot comparing the average gene expression for the top 50 differentially expressed genes in parasite cluster P0 for Caco-2 versus macrophage cells (left), Vero versus macrophage cells (middle) and Vero versus Caco-2 cells (right). **B.** Scatter plot comparing the average gene expression for the top 50 differentially expressed genes in parasite cluster P4 for Caco-2 versus macrophage cells (left), Vero versus macrophage cells (middle) and Vero versus Caco-2 cells (right). **C.** Scatter plot comparing the average gene expression for the top 50 differentially expressed genes in parasite cluster P5 for Caco-2 versus macrophage cells (left), Vero versus macrophage cells (middle) and Vero versus Caco-2 cells (right). **D.** Scatter plot comparing the average gene expression for the top 50 differentially expressed genes in parasite cluster P1 (left), P2 (middle), P3 (right) for Vero versus Caco-2 cells.

**Supplementary Figure 4: Transcriptional analysis of host response to infection in Caco-2 and Vero cells infected with *E. intestinalis*. A.** UMAP plot of host-only transcripts for all Caco-2 cells colored by the percentage of microsporidia transcripts (uninfected, gray and infected, purple) **B.** UMAP plot of host-only transcripts for all Vero cells colored by the percentage of microsporidia transcripts (uninfected, gray and infected, purple). **C.** Bar plot showing $-\log_{10}$ adjusted *p*-value from gene set enrichment analysis of biological pathways on host cluster C3 using the hallmark gene set in the Human Molecular Signatures Database (MSigDB) [31,32]. **D,E.** Quantification of the percentage of parasite developmental stages (P0-P5) found within the host clusters in Caco-2 **(D)** and Vero **(E)** infected cell datasets.

**Supplementary Table 1:** Differentially expressed parasite genes from each cluster from Caco-2 infected cell dataset.
**Supplementary Table 2:** Differentially expressed parasite genes from each cluster from Vero infected cell dataset.
**Supplementary Table 3:** Differentially expressed host genes from each cluster in the Caco-2 infected cell dataset (Host only transcriptome analysis).

**Supplementary Table 4:** Differentially expressed host genes from each cluster in the Vero infected cell dataset (Host only transcriptome analysis).
